# Evolution of specialization in heterogeneous environments: equilibrium between selection, mutation and migration

**DOI:** 10.1101/353458

**Authors:** Sepideh Mirrahimi, Sylvain Gandon

## Abstract

Adaptation in spatially heterogeneous environments results from the balance between local selection, mutation and migration. We study the interplay among these different evolutionary forces and demography in a classical two habitat scenario with asexual reproduction. We develop a new theoretical approach that fills a gap between the restrictive assumptions of Adaptive Dynamics and Quantitative Genetics. This analysis yields more accurate predictions of the equilibrium phenotypic distribution in different habitats. We examine the evolutionary equilibrium under general conditions where demography and selection may be non-symmetric between the two habitats. In particular we show how migration may increase differentiation in a source-sink scenario. We discuss the implications of these analytic results for the adaptation of organisms with large mutation rates such as RNA viruses.

## 1 Introduction

Spatially heterogeneous selection is ubiquitous and constitutes a potent evolutionary force that promotes the emergence and the maintenance of biodiversity. Spatial variation in selection can yield adaptation to local environmental conditions, however, other evolutionary forces like migration and mutation tend to homogenize the spatial patterns of differentiation and thus to impede the build up of local adaptation. Understanding the balance between these contrasted evolutionary forces is a major objective of evolutionary biology theory [Slatkin, 1978, Whitlock, 2015]). In this article, we consider a two-habitat model with explicit demographic dynamics as in [Meszéna et al., 1997, Day, 2000, Ronce and Kirkpatrick, 2001, Débarre et al., 2013]. We assume that adaptation is governed by a single quantitative trait where individuals reproduce asexually. Maladapted populations have a reduced growth rate and, consequently, lower population size. In other words, selection is assumed to be ‘hard’ [Christiansen, 1975, Débarre and Gandon, 2010] as the population size in each habitat is affected by selection, mutation and migration. These effects are complex because, for instance, asymmetric population sizes affect gene flow and adaptation feeds back on demography and population sizes [Nagylaki, 1978, Lenormand, 2002, Meszéna et al., 1997, Day, 2000, Ronce and Kirkpatrick, 2001, Débarre et al., 2013]. To capture the complexity of these feed backs it is essential to keep track of both the local densities and the distributions of phenotypes in each habitat. Note that this complexity often led to the analysis of the simplest ecological scenarios where the strength of selection, migration and demographic constraints are assumed to be the same in the two habitats (but see [Holt and Gaines, 1992, García-Ramos and Kirkpatrick, 1997, Gomulkiewicz et al., 1999, Holt et al., 2003] for analyses of asymmetric ecological scenarios). Three different approaches have been used to analyze this two-population model. Each of these approaches rely on a set of restrictive assumptions regarding the relative influence of the different evolutionary forces acting on the evolution of the population.

First, under the assumption that the rate of mutation is weak relative to selection it is possible to use the Adaptive Dynamics framework [Meszéna et al., 1997, Day, 2000, Débarre et al., 2013, Fabre et al.,]. This analysis captures the effect of migration and selection on the long-term evolutionary equilibrium. In particular, this approach shows that weak migration relative to selection promotes the coexistence of two specialist strategies (locally adapted on each habitat). In contrast, when migration is strong relative to selection, a single generalist strategy is favored. The main limit of this approach is that it relies on the assumption that there is a very limited amount of genetic variability. At most, 2 genotypes can coexist with these assumptions.

Second, Quantitative Genetics formalism has been used to track evolutionary dynamics in heterogeneous habitats when there is substantial level of phenotypic diversity in each population [Ronce and Kirkpatrick, 2001]. In this model the additive genetic variance is maintained by sexual reproduction but it may also be generated by large mutation rates in models with asexual reproduction [Débarre et al., 2013]. This formalism allows to recover classical migration thresholds below which specialization is feasible. But the analysis of [Ronce and Kirkpatrick, 2001] also reveals the existence of evolutionary bistability where transient perturbations of the demography can have long term evolutionary consequences on specialization. Yet, the assumption on the shape of the phenotypic distribution (assumed to be Gaussian in each habitat) is a major limit of this formalism.

Third, attempts to account for other shapes of the phenotypic distributions in heterogeneous environments have been developed recently [Yeaman and Guillaume, 2009, Débarre et al., 2013, Débarre et al., 2015]. These models highlight that calculations based on the Gaussian approximation which neglects the skewness of the equilibrium phenotypic distribution underestimates the level of phenotypic divergence and local adaptation. Yet, there is currently no model able to accurately describe the build up of non-Gaussian distributions. The only attempt to model this distribution is to describe the phenotypic distributions in each habitat as the sum of two Gaussian distributions [Yeaman and Guillaume, 2009, Débarre et al., 2013]. These models, however, only yield approximate predictions on long-term evolutionary equilibria.

Here we develop an alternative formalism that yields the phenotypic distribution in each habitat at the equilibrium between selection, mutation and migration. We start with the limiting case where the genetic variance due to mutations is very low, and we fully characterize the evolutionary equilibria of our system using Adaptive Dynamics. Second, we extend this analysis to a scenario where mutations are more frequent, and we derive approximations for the level of adaptation under a migration-selection-mutation balance. We also explore the effects of asymmetric constraints on selection, migration or demography between the two habitats. We evaluate the accuracy of these approximations by comparing them to numerical solutions of our deterministic model and we show that our approach improves previous attempts to study the interplay between adaptation and demography in heterogeneous environments. We contend that our results are particularly relevant for organisms with high mutation rates and may help to understand the within-host dynamics of chronic infections by RNA viruses [Drake and Holland, 1999, Sanjuán et al., 2010].

## 2 The model

We model an environment containing two habitats that we label 1 and 2 (Figure 1). The population is structured by a quantitative trait *z*. In each habitat there is selection towards an optimal value of the trait. Maladapted individuals suffer from a decreased growth rate: growth rate in habitat *i* is denoted by *r*_*i*_(*z*) which has its maximum *r*_max,*i*_ for the optimal traits *θ*_*i*_ which yields (for habitat *i* = 1,2):

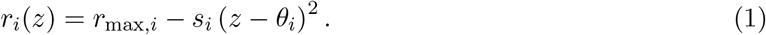

We denote by *s*_*i*_ the selection pressure in habitat *i*. Without loss of generality we assume that *θ*_1_ = −*θ*_2_ = −*θ*.

**Figure 1.**
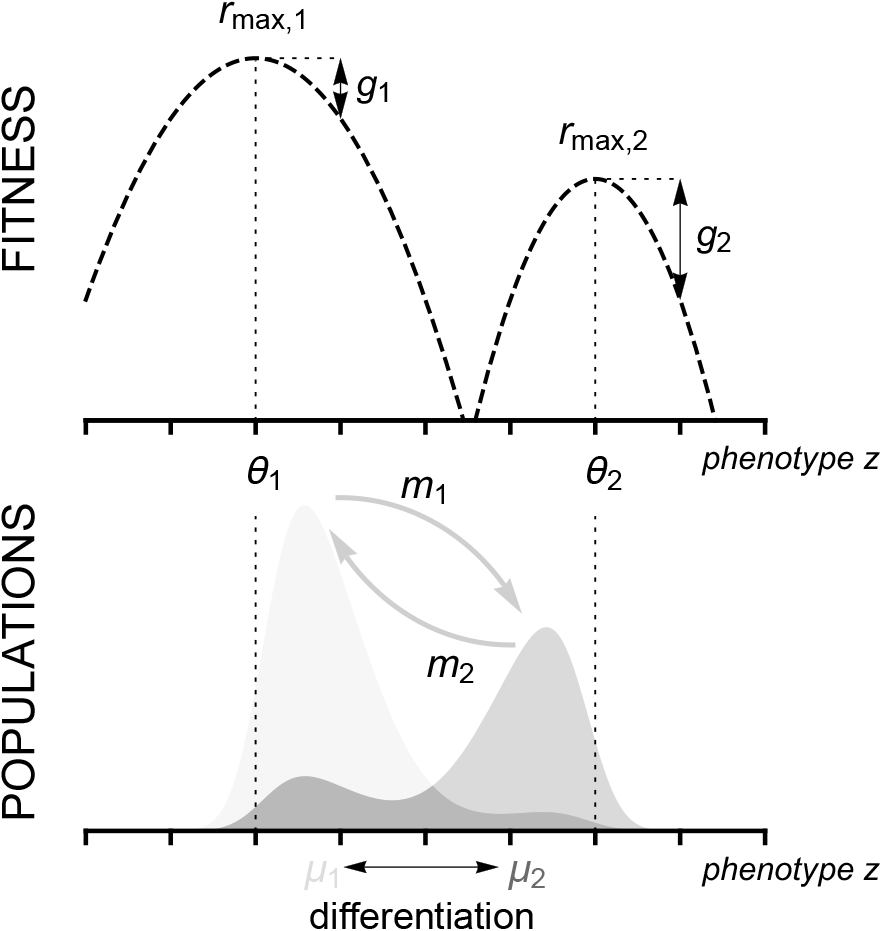
Schematic representation of the 2 habitat model. The top figure shows the growth rate (fitness) in each habitat as a function of the phenotypic trait *z*. In habitat *i* the growth rate is assumed to be maximized at *z* = *θ*_*i*_ and the strength of selection is governed by *s*_*i*_ (see equation (1)). Here we illustrate a scenario with asymmetric fitness functions. The bottom figure shows the phenotypic distribution in each habitat (light blue and light red in habitats 1 and 2, respectively). Migration from population *i* is governed by the parameter *m*_*i*_ and tends to reduce the differentiation (i.e. the difference between the mean phenotypes) between populations.

Reproduction is assumed to be asexual. Offsprings inherit the phenotype of their parent except when there is mutation (i.e. no environmental variance). We consider a continuum of alleles model [Kimura, 1965]. Mutation occurs with rate *U*, and add an increment *y* to the parents’ phenotype; we assume that the distribution of these mutational effects is given by *μ*(*y*), with mean 0 and variance equal to *V*. We also assume that individuals disperse among habitats with rates *m*_1_ and *m*_2_ which are independent of phenotype.

Let *n*_*i*_(*z*) be the phenotypic distribution in habitat *i* at time *t*. The dynamics of this distribution in each habitat is given by (for *i* = 1,2 and *j* = 2,1):

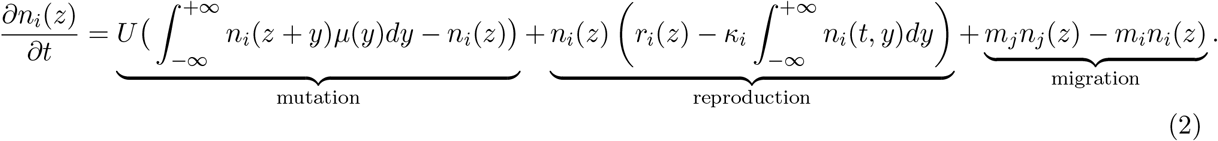

The first term in the right hand side of the above equation corresponds to the effect of mutations. The second term corresponds to logistic growth that results from the balance between reproduction given by (1) and density dependance where *κ*_*i*_ measures the intensity of competition within each habitat. The last term corresponds to the dispersal of individuals between habitats.

If we assume that the variance of the mutation distribution is small enough while the mutation rate *U* is not very small, we can consider an approximate model where we replace the mutation term in (2) by a diffusion (see [Kimura, 1965, Lande, 1975] and the more recent article [Champagnat et al., 2008] where the diffusion term has been derived directly from a stochastic individual based model). See also [Bürger, 2000]–pages 239-241 for a discussion on the domain of the validity of such model. Our model then becomes:

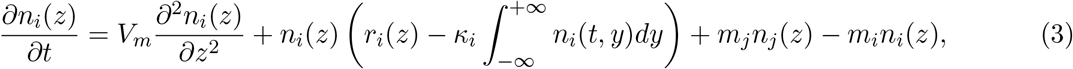

where *V*_*m*_ is proportional to *UV*. This model is close to the model studied in [Débarre et al., 2013]). The total population sizes in each habitat is given by:

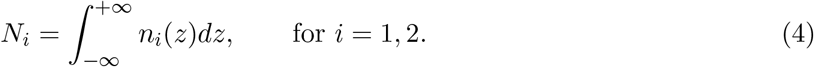

In other words, *n*_*i*_(*z*) refers to the *distribution of the phenotype* in habitat *i*, while *N*_*i*_ refers to the *density* of the polymorphic population in habitat *i*. Using (1) and (3) one can derive dynamical equations for the size of the population and the mean phenotype 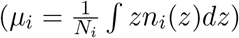:

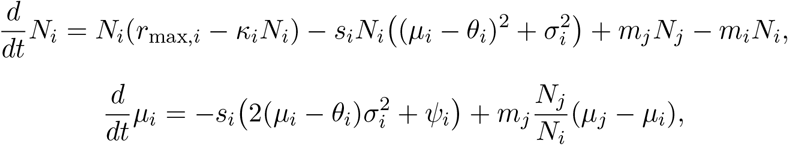

where 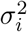 and *ψ*_*i*_ are the variance and the third central moment of the phenotypic distribution, respectively. These two quantities are also dynamical variables and their dynamics are governed by higher moments of the phenotypic distribution. These higher moments are also dynamical variables that depend on higher moments which indicates that we are dealing with a dynamical system that is not closed. Various approximations, however, have been used to capture its behavior. Typically, many results are based on the Gaussian approximation that focuses on the dynamics of the mean and the variance and discards all higher cumulants of the distribution [Bürger, 2000, Rice, 2004]. Yet several authors pointed out that neglecting the skewness of the distribution can underestimate the amount of differentiation and local adaptation [Yeaman and Guillaume, 2009, Débarre et al., 2013, Débarre et al., 2015].

Indeed, in the case of symmetric habitats, one can readily obtain the size and the mean trait of the population at equilibrium (the equilibrium is indicated by a superscript *). Using the fact that 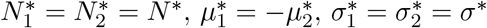 and 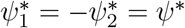, we obtain:

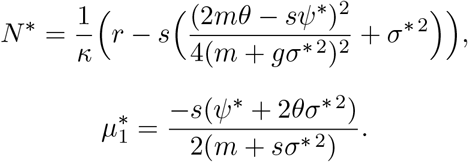

The differentiation between the two habitats is thus (Figure 1):

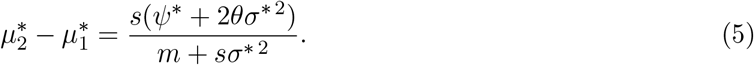

There is, however, no analytic predictions on the magnitude of the skewness of the phenotypic distribution except in the limit when the mutation rate is extremely low [Débarre et al., 2013].

## 3 The selection-mutation-migration equilibrium

We want to characterize the stationary solutions of (3) which results from the equilibrium between selection, mutation and migration in each habitat. In the following we present our two-step approach. First, we analyse the evolutionary equilibria of (3) when mutations are rare (i.e. *U* is vanishingly small). This allows to identify monomorphic or dimorphic evolutionary stable strategies (ESS). Second, we use these ESSs to derive an approximation for the stationary solutions of (3) when mutation is more frequent and maintains a standing variance at equilibrium.

### 3.1 Adaptive dynamics and evolutionary stable strategies

We consider a resident population at a demographic equilibrium set by the phenotypic distributions of the resident in both habitats. We want to determine the fate of a mutant with phenotype *z*_*m*_ introduced in this resident population. The ability of the mutant to invade is determined by its fitness given by (i.e. per capita growth rate minus density dependence):

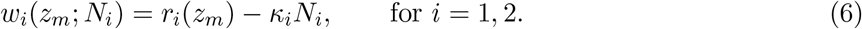

To take into account migration between habitats we introduce an effective fitness which corresponds to the growth rate of a trait in the whole environment (see [Szilágyi and Meszéna, 2009, Mirrahimi, 2013, Fabre et al.,]). The effective fitness *W*(*z*_*m*_; *N*_1_,*N*_2_), which corresponds to the *effective* growth rate associated with trait *z*_*m*_ in the resident population (*n*_1_,*n*_2_), is the largest eigenvalue of the following matrix:

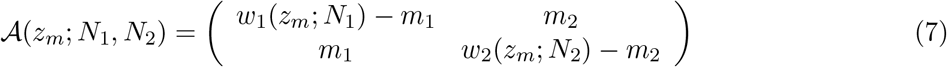

After some time the dynamical system will reach a stable demographic equilibrium. Because there are two habitats, we expect that at most two distinct traits can coexist. With an analysis of the effective fitness *W*, we characterize such equilibrium corresponding to the evolutionary stable strategy (see the supplementary information and [Mirrahimi, 2017]). This equilibrium is indeed either monomorphic (with phenotype *z*^*M*\*^ and the total population size 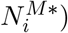 or dimorphic (with phenotypes 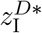 and 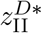 and the total population sizes 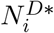, where the subscripts I and II indicate that the phenotype is best adapted to habitat 1 and 2, respectively).

### 3.2 Equilibrium distributions with mutation

In the above section (see also the supplementary information) we derived the evolutionary equilibria when mutations are very rare. These equilibria also correspond to a scenario where all the phenotypic strategies are present initially but where mutation is absent. In the following we allow mutation rate to increase and we study the impact of mutations on the ultimate evolutionary equilibrium of the phenotypic distributions. We present below the general principle of the approach before examining specific case studies.

We introduce a new parameter 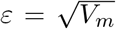 and hence in what follows we replace *V*_*m*_ by *ε*^2^, and we approximate the population’s phenotypical distribution *n*_*ε,i*_(*z*) in terms of *ε*.

Our method is based on the following ansatz:

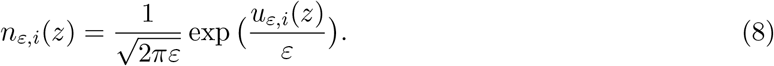

Note that a first approximation of the population’s distribution which is indeed commonly used in the theory of quantitative genetics is a Gaussian approximation of the following form around *z*\*:

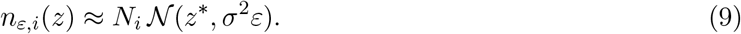

Another approximation that has been commonly used to study models of selection and mutation with continuous traits, is the house-of-cards approximation [Kingman, 1978, Turelli, 1984, Bürger, 2000]. This approximation is based on the assumption that the mutation distribution is independent of the original state of the gene. The house-of-cards approximation is valid in the limit of small mutation rates [Bürger, 2000]. Our analysis, however, is valid when the variance of the mutation distribution is small but the mutation rate is not vanishing. Our assumptions are thus closer to the domain of the validity of the Gaussian approximation. Yet, our objective is to obtain more accurate results than (9) and to approximate *u*_*ε,i*_ without making an a priori Gaussian assumption. To this end we postulate an expansion for *u*_*ε,i*_ in terms of *ε*:

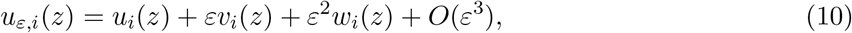

and we try to compute the coefficients *u*_*i*_(*z*), *v*_*i*_(*z*), and *w*_*i*_(*z*). First we can show that, when there is migration in both directions (i.e. *m*_*i*_ > 0 for *i* = 1,2), the zero order terms are the same in both habitats: *u*_1_(*z*) = *u*_2_(*z*) = *u*(*z*) (see the supplementary information). We can indeed compute explicitly *u*(*z*) which is given by (A.5) in the monomorphic case and by (A.6) in the dimorphic case. As we observe in the formula (A.5) and (A.6), *u*(*z*) attains its maximum (which is equal to 0) at the ESS points identified in the previous section. This means that the peaks of the population’s distribution are around the ESS points (*z*^*M*\*^ in the case of monomorphic ESS and 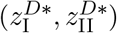 for the dimorphic ESS). We are also able to compute the first order term *v*_*i*_ and the value of *w*_*i*_ at the ESS points. This allows us to provide an approximation of the population’s density *N*_*ε,i*_ and the population’s distribution *n*_*ε,i*_ that we will call henceforth our *first approximation*. This approximation can be computed relatively easily (see [Mirrahimi, 2017]) and approximates very well the population’s distribution at equilibrium (see for instance Figure 2).

In order to provide more explicit formula for the moments of order *k* ≥ 1 of the population’s distribution in terms of the parameters of the model, we also provide a *second approximation*. This *second approximation,* instead of using the values of *u* and *v*_*i*_ in the whole domain, is based on the computation of a fourth order approximation of *u* and a second order approximation of *v*_*i*_ around the ESS points. Our *second approximation* is by definition less accurate than the first one. It still provides convincing results when the parameters are such that we are far from the transition zone from monomorphic to dimorphic distribution (see for instance Figure 2). This approximation is indeed based on an integral approximation which is relevant only when the population’s distribution is relatively sharp around the ESS points. This is not the case in the transition zone unless the effect of the mutations, i.e. *ε*, is very small.

## 4 Case studies

### 4.1 Symmetric fitness landscapes

We focus first on a symmetric scenario where, apart from the position of the optimum, the two habitats are identical: *m*_1_ = *m*_2_ = *m*,*k*_1_ = *k*_2_ = *k*, *s*_1_ = *s*_2_ = *s*,*r*_max,1_ = *r*_max,2_ = *r*_max_. In this special case it is possible to fully characterize the evolutionary equilibrium.

When migration rate is higher than critical migration threshold *m* > *m*_*c*_ = 2*sθ*^2^ migration prevents the differentiation of the trait between the two habitats [Mirrahimi, 2017]. The evolutionary equilibrium, when the mutation rate is vanishingly small, is monomorphic and satisfies *z*^*M*\*^ = 0 and 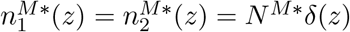 where *δ*(.) is the dirac delta function and 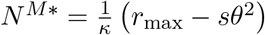.

#### Monomorphic case

Let’s suppose that *m*_*c*_ = 2*sθ*^2^ ≤ *m*. Then *z*^*M*\*^ = 0 is the only ESS and 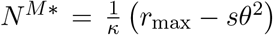. Then, the population’s distribution *n*_*ε,i*_(*z*) can be approximated following the method introduced above. In particular, defining 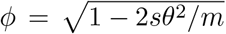 we can approximate the moments of the population’s distribution (Figure 2):

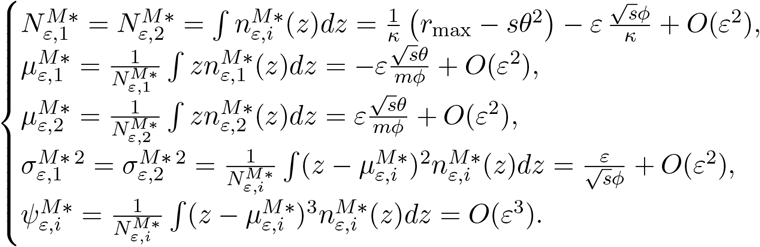

Note that the variance 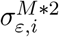 is larger than the case with no heterogeneity between the habitats, where we recover the well-known equilibrium variance 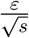 of quantitative genetics [Lande, 1975, Bürger, 2000, Rice, 2004]. This increase of the variance comes from *ϕ* which depends on dispersion and the heterogeneity between the two habitats. The variance of the distribution increases as *ϕ* decreases. When *ϕ* = 0 the approximation for the variance becomes infinitely large. This corresponds to the threshold value of migration below which the above approximation collapses because the distribution becomes bimodal. In this case we have to switch to the analysis of the dimorphic case. Note also that the differentiation between habitats depends also on *ϕ*. Some differentiation start to emerge even when the migration rate is above the critical migration rate, *m*_*c*_ (Figures 2 and 3).

**Figure 2.**
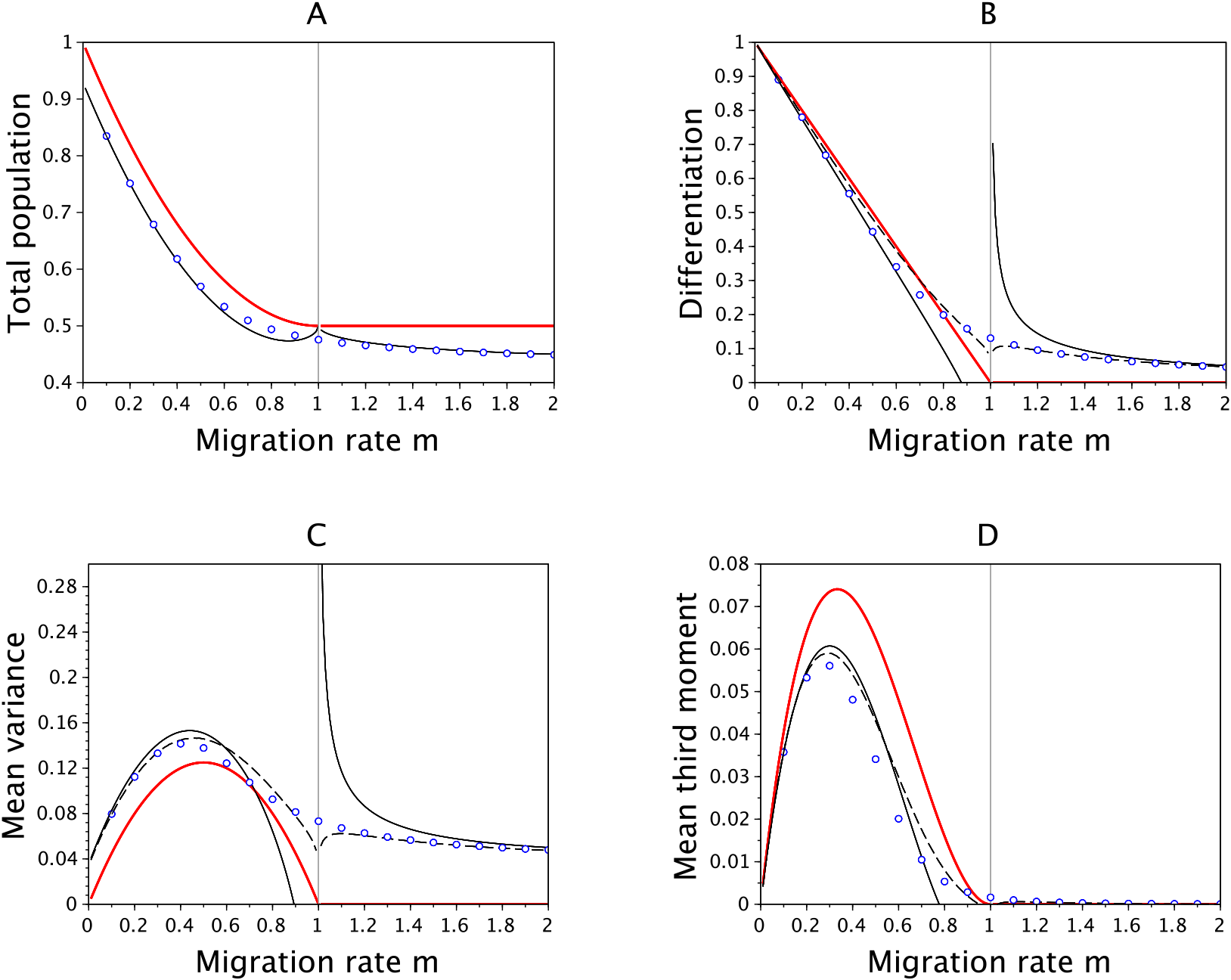
Effects of migration in a symmetric scenario on (A) the total population size in habitat 1, (b) the differentiation between habitats, (c) the variance and (d) the third central moment of the phenotypic distribution in habitat 1. The dots refer to the numerical resolutions of the problem with *ε* = 0.05, the red line indicates the case where *ε* = 0 while the lines in black refer to our two approximations when *ε* = 0.05 (the dashed line for the *first approximation* and the full line for the *second approximation*). The vertical gray line indicates the critical migration rate below which dimorphism can evolve in the adaptive dynamics scenario. Other parameter values: *r*_max_ = 1, *s* = 2, *θ* = 0.5, *κ* = 1. Note that, in this figure and in the following ones, to compute numerically the equilibrium, we have solved numerically the dynamic problem (3) and kept the solution obtained after long time when the equilibrium has been reached.

#### Dimorphic case

When *m* < *m*_*c*_, the only globally stable evolutionary equilibrium is dimorphic which yields the following ESS: 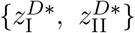 with 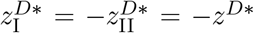 and 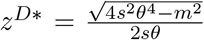. This yields the following equilibrium distribution when *ε* = 0: 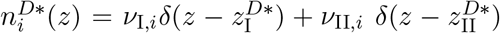 (analytic expressions for 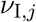 and 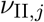 are given in the supplementary information). One can then approximate the local moments of the population’s distribution *n*_*ε,i*_(*z*), with 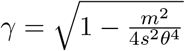,

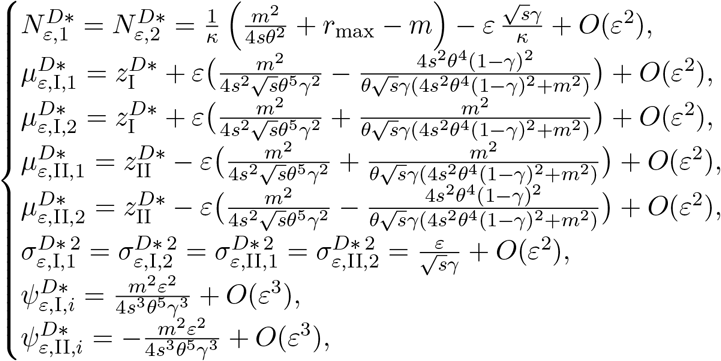

where the subscripts I and II indicate that the local moments are in the sets 𝒪_I_ = (−∞,0) and 𝒪_II_ = (0,∞) which include respectively 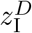 and 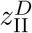 (see the supplementary information for the precise definitions). One can compute the global moments of the population’s distribution from the above local moments.

**Figure 3.**
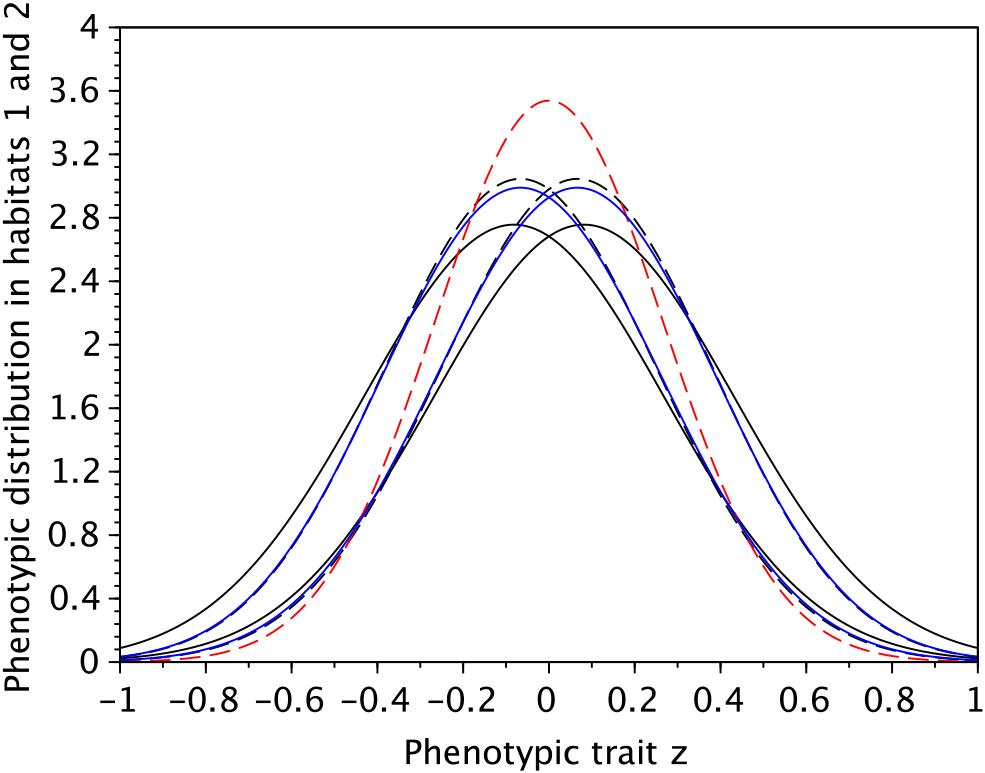
Selection-mutation-migration equilibrium distributions in the two habitats in a symmetric scenario and in a case where the population’s distribution is unimodal in each habitat. We plot the exact equilibrium distributions obtained from numerical computations (blue line) together with our 2 approximations (dashed lines for the *first approximation* and full black line for the *second approximation*) and the approximation given in [Débarre et al., 2013] (red line). Note that our approximations capture the emergence of some differentiation even though we are above the critical migration rate leading to the evolution of a dimorphic population. In presence of mutations, the population’s distribution is indeed shifted to the left(respectively right) in the first(respectively second) habitat, while [Débarre et al., 2013] provided the same approximation for both habitats. Note also that our approximation yields better approximations for the variance of the distribution in each habitat ([Débarre et al., 2013] underestimates this variance). Parameter values: *m* = 1.5, *r*_max_ = 3, *s* = 2; *θ* = 0.5, *κ* = 1, *ε* = 0.1.

### 4.2 Non-symmetric scenarios

#### A general non-symmetric scenario

In a non-symmetric scenario there also exists a unique ESS which is either monomorphic or dimorphic. There is still a threshold value of migration above which the maintenance of a dimorphic polymorphism is impossible: 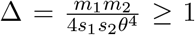. Note that this condition generalizes the condition in the symmetric case (i.e. when *m*_1_ = *m*_2_ and *s*_1_ = *s*_2_). However, for the ESS to be dimorphic, the condition Δ < 1 is not enough and two other conditions should also be satisfied. These conditions (i.e. *η*_1_ < *β*_2_*r*_max,2_ – *α*_1_*r*_max,1_ and *η*_2_ < *β*_1_*r*_max,1_ – *α*_2_*r*_max,2_ with the constants *α*_*i*_,*β*_*i*_ and *η*_*i*_ depending on the parameters *m*_1_, *m*_2_, *s*_1_, *s*_2_, *k*_1_, *k*_2_ and *θ*, see the supplementary information), guarantee that the qualities of the habitats are not very different. Indeed, if one habitat has a higher quality it is likely to overwhelm the dynamics of adaptation to the other habitat. This will yield a monomorphic equilibrium biased toward the high quality habitat. Figure 4 illustrates that a polymorphism is only maintained in a range of parameter values where the two habitats are relatively similar. Interestingly, in spite of the asymmetry of the two habitats, the locations of the two peaks of the phenotypic distribution are always symmetric and consequently: 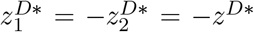 where: 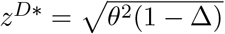. See the supplementary information for the expressions of the densities in each habitat and [Mirrahimi, 2017] for the derivation of this stable equilibrium.

#### A source-sink scenario

An extreme case of asymmetry occurs when one population (the source) does not receive any migrant from the second population (the sink). For instance, when *m*_1_ > 0 and *m*_2_ = 0 there is no immigration in habitat 1. Note that, this is a degenerate case and in particular, we are not anymore in the framework of Section 3.1, where the ESS was always the same in the both habitats as a result of strict positivity of migration rate in both directions. Moreover, the computation of the equilibrium in presence of mutations is also slightly different because of this degeneracy (see [Mirrahimi, 2017] for more details). In particular, we are not able to compute the first order corrector *v*_2_ in the whole domain. However, a local approximation of *v*_2_ gives already convincing results (Figure 5).

The evolutionary outcome in the first habitat is obvious because it depends only on selection acting in habitat 1: the ESS is −*θ* and

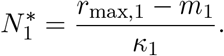

Moreover, the population’s phenotypically distribution *n*_*ε*,1_ can be computed explicitly: *n*_*ε*,1_ = *N*_*ε*,1_*f*_*ε*_, where 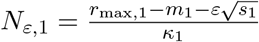 and *f*_*ε*_ is the probability density of a normal distribution 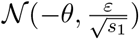.

In habitat 2, the evolutionary outcome results from the balance between migration from habitat 1 and local selection and two situations can arise (Figure 5): (i) monomorphic case: under the condition

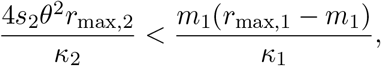

the evolutionary equilibrium is monomorphic and the evolutionary stable strategy is *z*\* = −*θ*. And the total population is given by

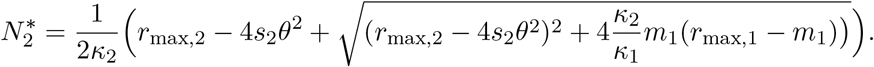

There is indeed a population of size 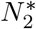 in the second habitat which is of type *z*\* = −*θ*: this population is very maladapted.

**Figure 4.**
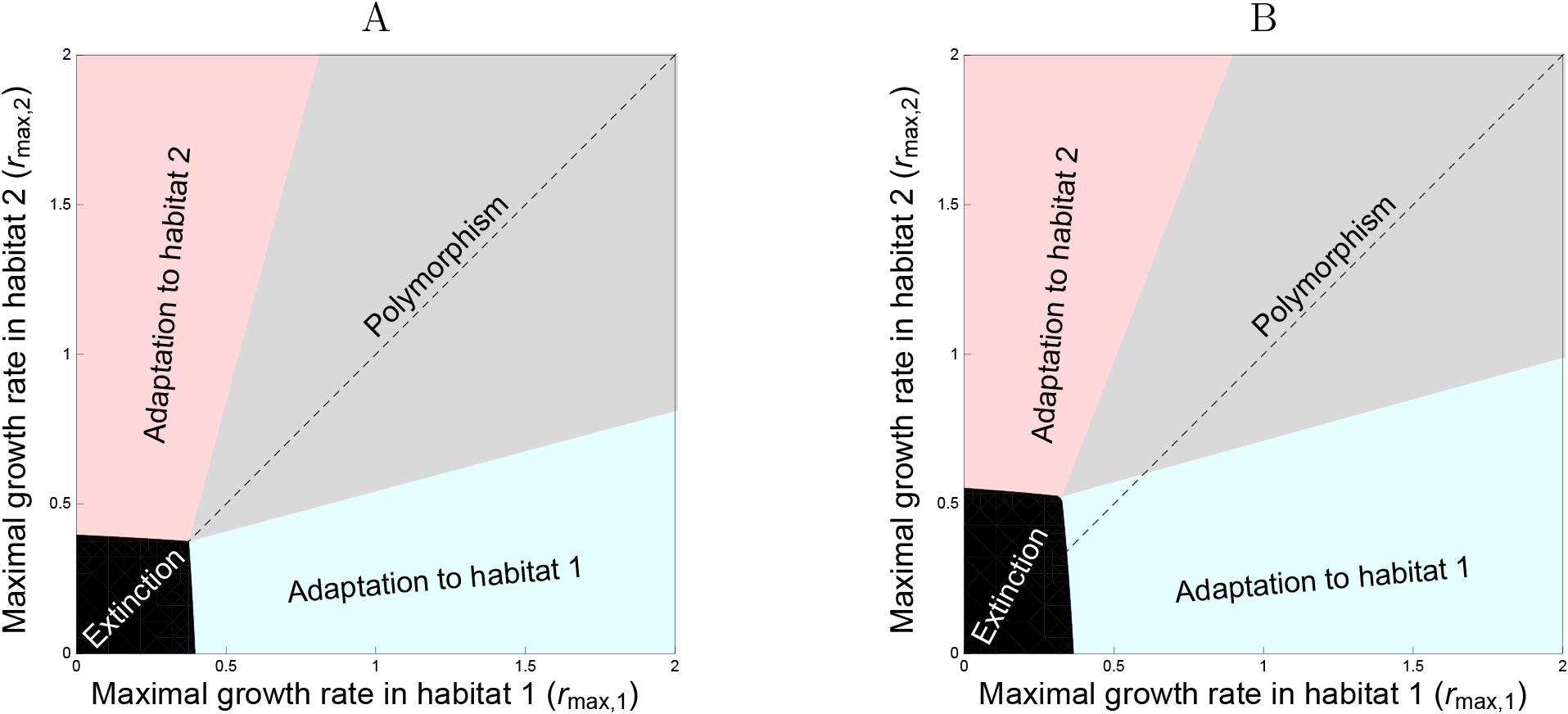
Maintenance of polymorphism and asymmetric adaptation as a function of the maximal growth rates *r*_max,1_ and *r*_max,2_ in the two habitats. In (A) we examine a symmetric situation where all the parameters are identical in the two habitats: *m*_1_ = *m*_2_ = 0.5, *s*_1_ = *s*_2_ = 2, *k*_1_ = *k*_2_ = 1. In (B) we show an asymmetric case with the same parameters as in (A) except *m*_1_ = 0.5 and *m*_2_ = 0.7. The black area indicates the parameter space where the population is driven to extinction because the maximal growth rates are too low. In the grey area some polymorphism can be maintained in the two-habitat population as long as the difference in the maximal growth rates are not too high. When this difference reaches a threshold polymorphism cannot be maintained and the single type that is maintained is more adapted to the good-quality habitat (the habitat with the highest maximal growth rate).

The population’s distribution *n*_*ε*,2_(*z*) can also be approximated. In particular we can approximate the moments of the population’s distribution:

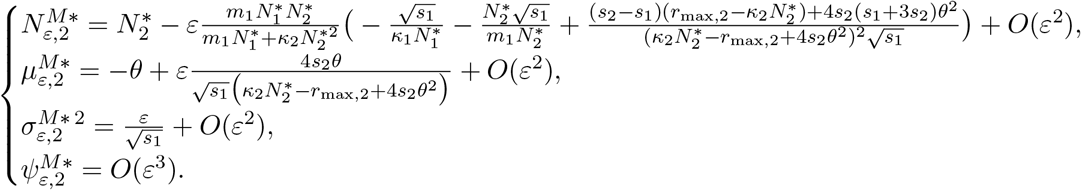

When the above condition is not satisfied, i.e.

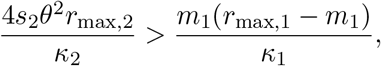

the evolutionary equilibrium is dimorphic in the second habitat (Figure 5):

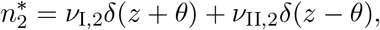

with

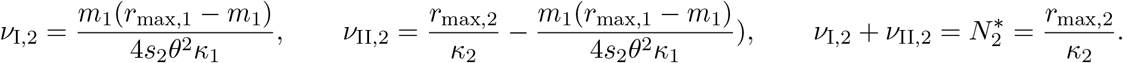

Then, the moments of the population’s distribution *n*_*ε*,2_(*z*) can be approximated as below:

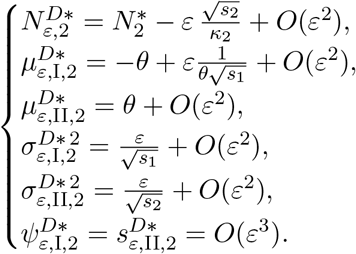

This source-sink scenario illustrates the complex and unexpected effects of migration when there is an asymmetry between the two habitats. Indeed, Figure 5A shows that the population size in the sink is maximized for intermediate values of migration. More migration from the source has a beneficial effect on the demography of the sink but it prevents local adaptation. Yet, when migration from the source becomes very strong it limits the growth rate of the source. This limits the influence of the source on the sink and may even promote adaptation to the sink. In fact it is worth noting that differentiation between the two habitats can actually increase with migration (Figure 5B).

## 5 Discussion

We derive approximations for the equilibrium distribution between selection, mutation and migration in a two-habitat environment. The derivation starts with the analysis of our model when the mutation rate is assumed to be vanishingly small. This Adaptive Dynamics approach allows to characterize the long-term evolutionary outcome. First, when migration is strong relative to selection, the population generally evolves towards a monomorphic evolutionary stable equilibrium. Second, when migration is limited the population generally evolves towards a dimorphic evolutionary equilibrium. Our derivation in the small mutation limit generalizes the results obtained in previous studies [Ronce and Kirkpatrick, 2001, Débarre et al., 2013] to scenarios where the two habitats may be asymmetric. In particular we show that the condition for the maintenance of a two specialized strategies are more restrictive with asymmetric scenarios (Figure 4). Indeed, asymmetries promote a single strategy that is more locally adapted to the habitat with larger population size and/or lower immigration rate. The fact that asymmetric migration promotes monomorphism was also observed in a related model [Akerman and Bürger, 2014]. The analysis of an extreme case with source-sink dynamics reveals the complex interplay between migration, demography and local selection. The maintenance of a polymorphic equilibrium is possible when migration from the source is either very weak or very strong. This result challenges the classical prediction where migration is always an homogenizing force reducing the differentiation among populations (Figure 5).

Our approach allows to derive approximations for equilibrium phenotypic distributions under larger mutation rates. In the symmetric scenario we recover the classical results from quantitative genetics [Lande, 1975, Bürger, 2000, Rice, 2004] but expand this to heterogeneous scenarios. In particular, we capture the emergence of differentiation between habitats when the migration rate decreases. When migration is strong relative to selection, the stationary distribution is weakly affected by spatially heterogeneous selection. When migration is close to the critical migration rate *m*_*c*_ we predict the build up of some differentiation and the maintenance of a higher amount of variation in each habitat ( Figure 3). When the migration rate is much smaller than *m*_*c*_ selection is sufficiently strong between habitats and the equilibrium distribution in each habitat can be well approximated as the sum of two distributions. But unlike previous approximations [Yeaman and Guillaume, 2009, Débarre et al., 2013] these two distributions are non-gaussian. We derive approximations for the moments of these distributions. In other words, this work generalises previous attempts to derive the distribution of a phenotypic trait at the mutation-selection-migration equilibrium. Our results confirm the importance of the skewness in the phenotypic distribution and improve predictions of measures of local adaptation in a heterogeneous environment.

**Figure 5.**
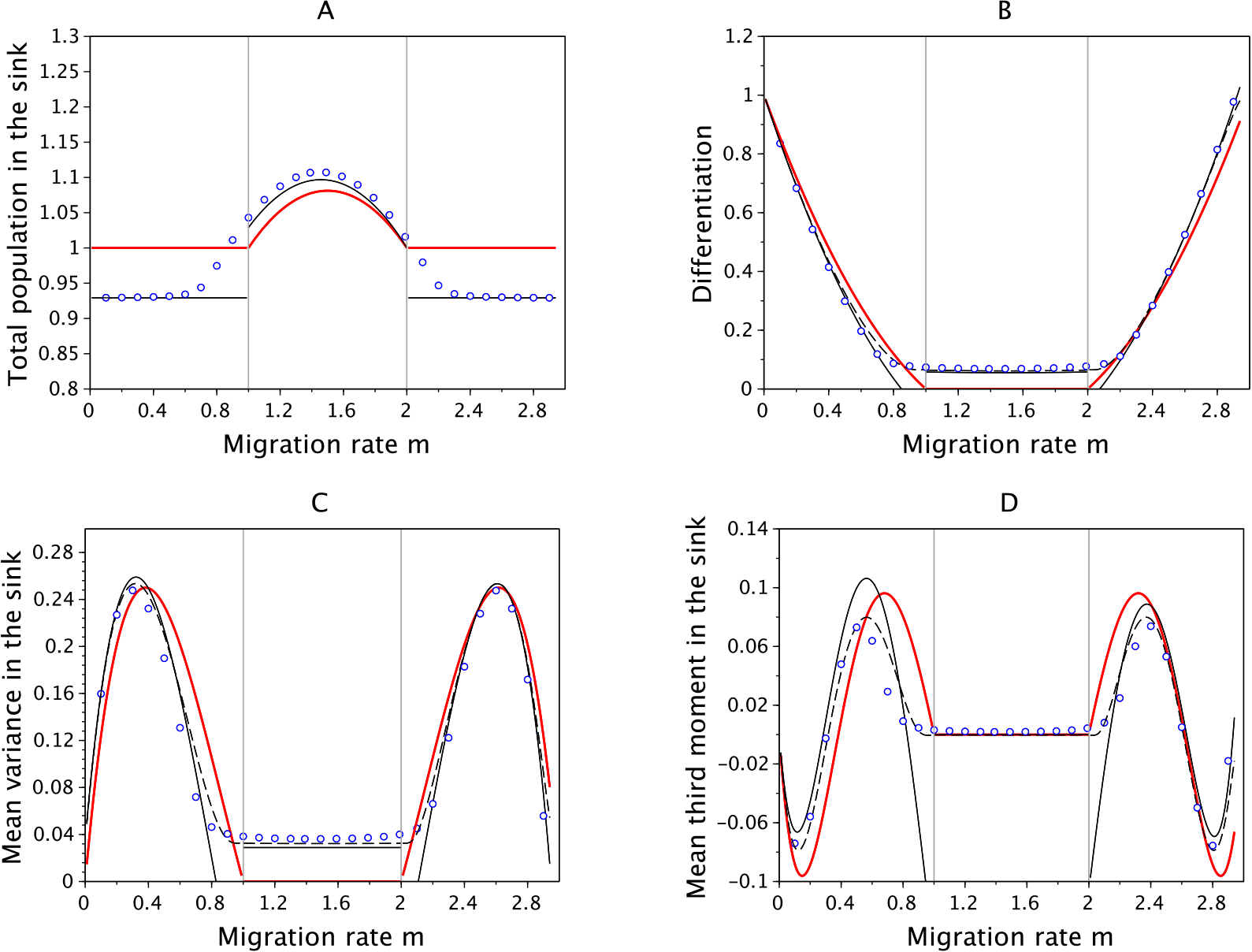
Effects of migration in a source-sink scenario on (A) the total population size in the sink habitat, (B) the differentiation between habitats, (C) the variance and (D) the third central moment of the phenotypic distribution in sink. The dots refer to exact numerical computations when *ε* = 0.05, the red line indicates the case where *ε* = 0 while the lines in black refer to our two approximations when *ε* = 0.05 (dashed line for the *first approximation* and the full line for the *second approximation*). The vertical gray line indicates the critical migration rates where transition occurs from monomorphism to dimorphism in the adaptive dynamics framework. Other parameter values: *r*_max,1_ = 3, *r*_max,2_ = 1, *s*_1_ =3, *s*_2_ = 2, *k*_1_ = *k*_2_ = 1, *θ* = 0.5.

Our work illustrates the potential of a new mathematical tool in the field of evolutionary biology.

In this work, we use an approach based on Hamilton-Jacobi equations (see (A.4)) which has been developed, mostly by the mathematical community, during the last decade to describe the asymptotic solutions of the selection-mutation models, as the effects of the mutations vanish. We refer to [Diekmann et al., 2005, Perthame and Barles, 2008, Mirrahimi, 2011] for the establishment of the basis of this approach. Note, however, that previous studies were mainly focused on the limit case where the effects of mutations are vanishingly small. In the present work we go further than the previous studies and characterize the whole phenotypic distributions when the effect of mutation is more important. Understanding the build up of this distribution is particularly important to study the effect of mutation on adaptation. Although mutation is the ultimate source of adaptive variation, the accumulation of deleterious mutations may also generate a load on the average fitness of populations. This is particularly relevant in organisms like RNA viruses which are characterized by very large mutation rates [Drake and Holland, 1999, Sanjuán et al., 2010]. In fact, the mutation loads of RNA virus is so high that it may even lead some populations to extinction [Bull et al., 2007, Martin and Gandon, 2010]. Our model can be used to accurately capture the effect of increasing mutation rates on the mutation load of a population living in a heterogeneous environment (Figure 6). This heterogeneity may be particularly relevant in chronic infections by pathogenic virus that can adapt to different organs [Kemal et al., 2003, Sanjuán et al., 2004, Ducoulombier et al., 2004, Jridi et al., 2006]. A better understanding of the equilibrium phenotypic distribution in heterogeneous environments may thus provide more accurate prediction on the critical mutation rates that can ultimately lead within-host dynamics to pathogen extinction.

**Figure 6.**
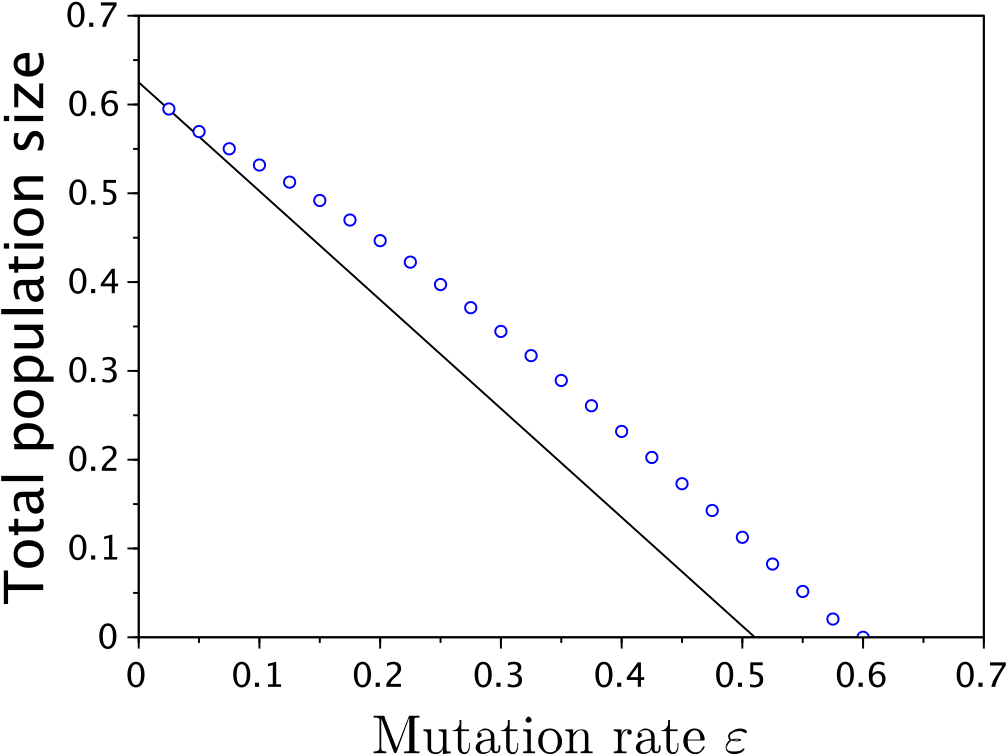
Effect of increasing the mutation rate *ε* on the total population size in the symmetric scenario used in figure 2 with *m* = 0.5. The full line indicates the approximation and the dots are the results of exact numerical computations. This figure illustrates that our approximation captures reasonably well the effect of large mutation rates on the mutation load in a two-habitat scenario where there is differentiation and some local adaptation.

Our analysis of the equilibrium between selection, migration and mutation could be extended in several new directions. More than 2 habitats could be considered, or different growth rates and/or mutation kernels could be used (see [Mirrahimi, 2013] and the supplementary information). The approach could also be used to analyze situations away from the equilibrium. For instance, it would be possible to track the dynamics of the distribution as the population adapts to a new environment or to a time-varying environment [Lande and Shannon, 1996]. Hamilton-Jacobi equations have indeed also been used to study time-varying (but space homogeneous) environments (see for instance [Mirrahimi et al., 2015]). Finally, it is interesting to note that the generalization of the present ecological scenario to model the adaptation of sexual species in heterogeneous environments remains to be carried out.

## Acknowledgements

The first author is grateful for partial funding from the European Research Council (ERC) under the European Union’s Horizon 2020 research and innovation programme (grant agreement No 639638), held by Vincent Calvez, and from the french ANR projects KIBORD ANR-13-BS01-0004 and MOD-EVOL ANR-13-JS01-0009.

## Supporting Information

### A Mathematical derivation

#### A.1 Adaptive dynamics

In this section, we provide the conditions for an evolutionary stable strategy. To be able to characterize the ESS one should first characterize the demographic equilibrium corresponding to a set of traits. Because there are only two habitats, we expect that at most two distinct traits can co-exist. Therefore, we only need to consider two scenarios where the phenotypic distribution is either monomorphic (with phenotype *z*^*M*^) or dimorphic (with phenotypes 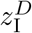 and 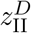, where the subscripts I and II indicate that the phenotype is best adapted to habitat 1 and 2, respectively).

The monomorphic equilibrium is given by 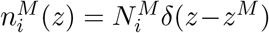 where *δ*(.) is the dirac delta function, 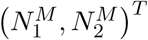 is the right eigenvector associated with the dominant eigenvalue 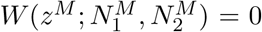 of 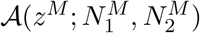. In a similar way the dimorphic equilibrium is characterized by: 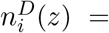 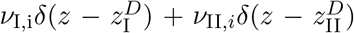, where 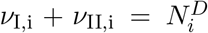 and (*v*_*k*,1_,*v*_*k*,2_)^T^ are the right eigenvectors associated with the largest eigenvalues 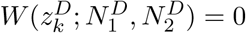 (for *k* = I, II) of 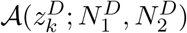.

The evolutionary stability of a resident strategy *z*^*M*\*^ can be studied with the analysis of the invasion of a new mutant strategy *z*_*m*_ at the demographic equilibrium (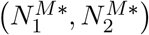 set by the resident strategy. The monomorphic strategy *z*^*M*\*^ is an evolutionary stable strategy if for any mutant *z*_*m*_ ≠ *z*^*M*\*^, the effective fitness is negative: 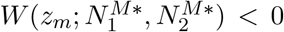. In a similar way, the dimorphic strategy 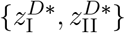 is an evolutionary stable strategy if for any mutant 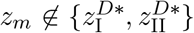, the effective fitness is negative: 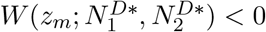.

See [Mirrahimi, 2017] where such evolutionary stable strategies are characterized.

#### A.2 Derivation of our *first approximation*

Our *first approximation* is based on the computation of the terms *u*_*i*_ and *υ*_*i*_ and the value of *w*_*i*_ at the ESS points. Based on such computations we can provide an approximation of the population’s density *N*_*ε*,i_ and the population’s distribution *n*_*ε*,i_ in the following form

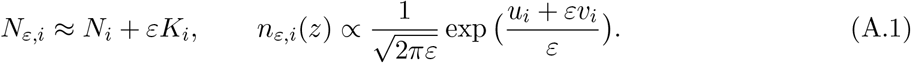

Indeed we neglect the next order terms in our approximation since when *ε* is small, in view of (8), they have only small contribution to the population’s distribution.

Note that the equilibrium (*n*_*ε*,1_,*n*_*ε*,2_) solves

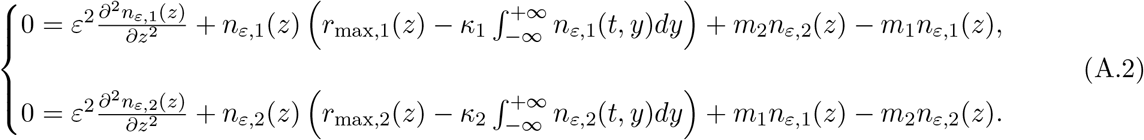

Replacing (8) in the above equation we obtain

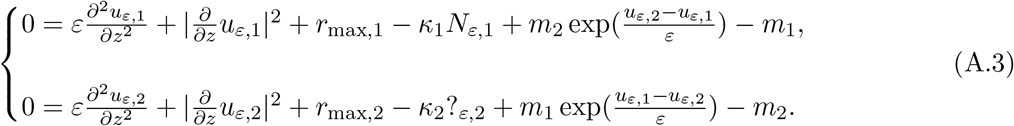

We can determine *u*_*i*_, *v*_*i*_ and the value of *w*_*i*_ at the ESS points from the above equation and (10). We show here how to obtain *u*_*i*_ (see [Mirrahimi, 2017] for more details and the computation of *v*_*i*_ and the value of *w*_*i*_ at the ESS points).

Note that the exponential terms in (A.3) suggest that, when *m*_*i*_ > 0 for *i* = 1,2, as *ε* → 0 *u*_*ε*,1_ and *u*_*ε*,2_ converge to the same limit *u*. One can prove indeed that *u*_*ε*,1_ and *u*_*ε*,2_ converge to a solution of the following Hamilton-Jacobi equation

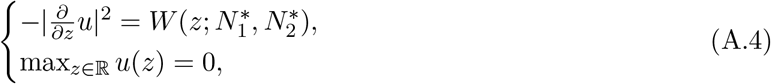

where 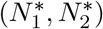 is the demographic equilibrium corresponding to the ESS and *W*(*z*; *N*_1_, *N*_2_), which is nonpositive, is the largest eigenvalue of the following matrix:

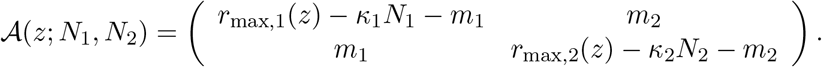

Note that *W* is indeed the effective fitness introduced in Section 3.1. One can also verify that (A.4) implies

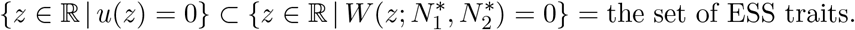

This property, together with the fact that there exists a unique ESS for this model [Mirrahimi, 2017], implies that the solution of (A.4) is unique.

In the case of monomorphic ESS, *u* is given by

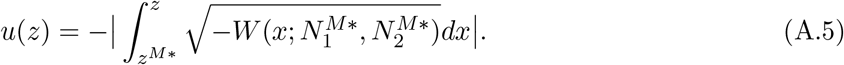

One can verify that *u*, given by the formula above, is smooth and solves (A.4) with its maximum point at *z*^*M*\*^. Note that the absolute values are necessary since the upper limit of the integral *z* can be smaller or larger than the lower limit *z*^*M*\*^.

In the case of dimorphic ESS, *u* is given by

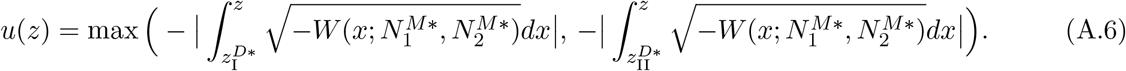

One can also verify that the above function is smooth at all points except at the point where the two functions in the maximum operator intersect. Moreover, *u* solves (A.4) at the smooth points and it attains its maximum at the ESS points 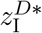 and 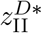. See [Mirrahimi, 2017] for the details on why this is indeed the solution obtained as the limit *ε* → 0.

#### A.3 Derivation of our *second approximation* and *N*_*ε,i*_

In this section, we provide the main idea to obtain explicit formula for the moments of the population’s distribution, and we precise the difference between our two approximations.

The computation of explicit formula for the moments of the population’s distribution is based on the observation that, when *ε* is small, the population’s distribution *n*_*ε,i*_(*z*) is exponentially small far from the ESS points, since *u*(*z*) takes negative values at those points. Therefore, only the values of *u*, *v*_*i*_ and *w*_*i*_ around the ESS points matter. We can show that to obtain a good approximation of the population’s distribution, with an error of order *ε*^2^ on the moments, it is enough to compute a fourth order approximation of *u*, a second order approximation of *v*_*i*_ and a zero order approximation of *w*_*i*_ around the ESS points.

We first provide our analytic formula for the moments of the population’s distribution in the monomorphic and dimorphic cases. We next show how to compute such approximations, in the case where the ESS is monomorphic.

##### A.3.1 Analytic formula for the moments of the population’s distribution

###### Monomorphic case

Let’s suppose that the model has a monomorphic ESS *z*^*M*\*^. In order to provide an explicit approximation of the moments of the population’s distribution, we compute the fourth order approximation of *u*(*z*) around *z*^*M*\*^:

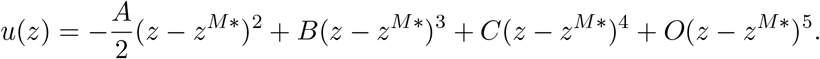

and the second order approximation of *v*_*i*_(*z*) around *z*^*M*\*^:

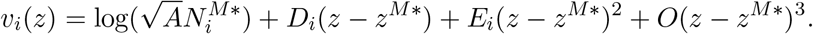

We also denote

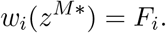

See [Mirrahimi, 2017] for the details of the computations of the above coefficients.

The above approximation allows us to estimate the moments of the population’s distribution:

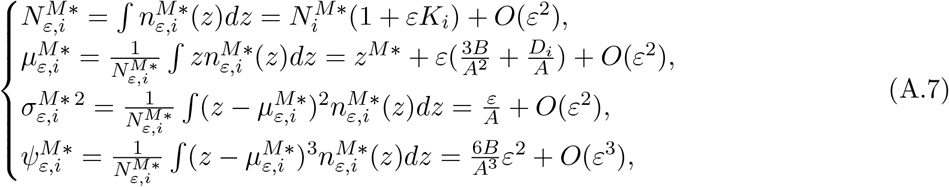

with

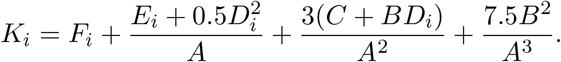

We notice that, even in this monomorphic case, because of the terms *D*_*i*_, the mean trait in the habitats are slightly different and there is a shift between the phenotypical distributions of the habitats. The variance, up to order *ε*, is determined knowing only the constant *A*. Finally the last line indicates that there is a non-zero skewness in the distribution when the constant *B* is nonzero, which may arise in a non-symmetric scenario.

###### Dimorphic case

Let’s suppose that the model has a dimorphic ESS 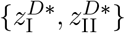 with 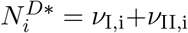. Similarly to above, we will use a Taylor expansion of *u*(*z*) around 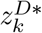, which can be easily computed from the above formula:

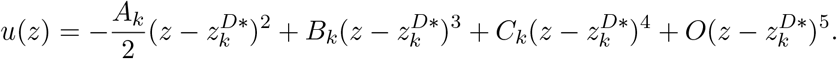

Following similar computations as in the case of monomorphic ESS one can also compute a second order approximation of exp(*v*_*i*_(*z*)) around 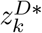 and the value of 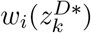:

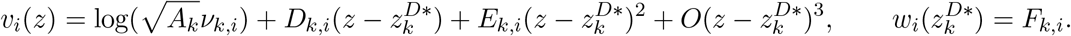

The above approximation allows us to estimate the local moments of the population’s distribution:

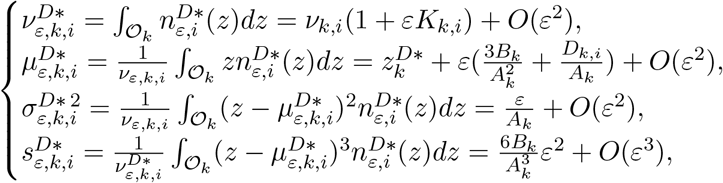

with

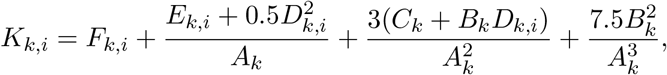

and 𝒪_I_ = (−∞,0) and 𝒪_II_ = (0,∞). Note also that one can compute the global moments of the population’s distribution from the above local moments.

##### A.3.2 Derivation of the analytic formula

We next show how to compute such approximations, in the case where the ESS is monomorphic. The computations for the dimorphic case follow similar arguments. To compute our approximations, we use the asymptotic expansion of *u*_*ε,i*_:

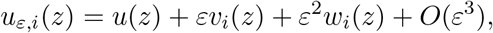

and the Taylor expansions of *u*, *v*_*i*_ and *w*_*i*_:

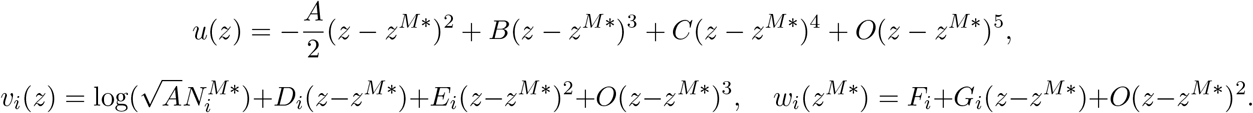

To obtain the zero order term in the expansion for *v*_*i*_(*z*) we use the fact that, as the mutation’s variance vanishes (*ε* → 0), the total population size *N*_*ε,i*_ tends to 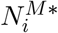 which corresponds to the demographic equilibrium at the ESS.

One can indeed use the above expressions to compute

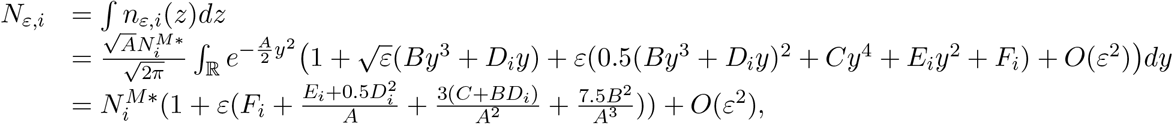

and for any integer *k* ≥ 1,

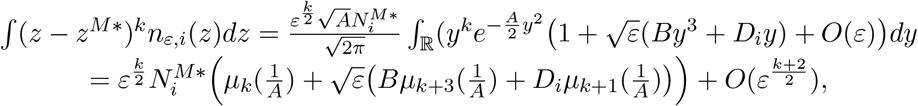

where *μ*_*k*_(*σ*^2^) corresponds to the k-th order central moment of a Gaussian distribution with variance *σ*^2^. Note that to compute the integral terms above we have performed a change of variable 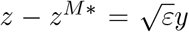, therefore each term *z* – *z*^*M*\*^ can be considered as of order 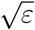 in the integrations. The above integrations are the main ingredients to obtain the approximations given in (A.7), i.e. our *second approximation*.

Note that the approximation of the total population size *N*_*ε,i*_ given above is also the one used in our *first approximation*. One can indeed observe from the above formula that in the computation of the total population size, in the contrary to the computation of the next order moments, there is a contribution of the second order term *w*_*i*_, via it’s value at the ESS point *F*_*i*_. This is why to obtain an approximation of *N*_*ε,i*_ with an error of order *ε*^2^ in our *first approximation,* it is not enough to only use the functions *u* and *v*_*i*_, which are computed globally, and the local value of *w*_*i*_, is still needed. However, to compute the mean moments of higher order in our *first approximation*, we perform a numerical integral using the approximation of *n*_*ε*_ given in (A.1) with the global values of *u* and *v*_*i*_, therefore no local approximation is made (except for the sink and source case which is a degenerate case).

#### A.4 Derivation of a Hamilton-Jacobi equation in the case of model (2)

Our approach can also be used to study the more general model (2). The objective would be to provide an approximation of the solution *n*_*i*_, when the variance of the mutation distribution is small. We assume indeed that the variance of the mutation distribution *μ*_*ε*_ scales as *ε*^2^*V*_o_. More precisely, we assume that 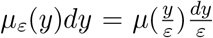 (for instance a Gaussian distribution with Variance *ε*^2^*σ* has such form). Then, the stationary version of (2) may be written as

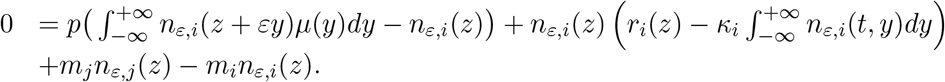

Next, analogously to our work in the case of (3), we use the ansatz (8):

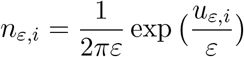

and postulate an expansion for *u*_*ε,i*_ in terms of *ε*:

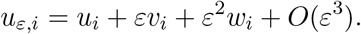

The computation of the above terms allows us to provide approximations of the population distributions *n*_*ε,i*_ and their moments. To compute these terms, analogously to what we presented for the diffusion case, thanks to the combination of the above equalities, we derive some equations satisfied by *u*_*i*_, *v*_*i*_ and *w*_*i*_. The resolution of such equations, which is less straight forward comparing to the diffusion case, allows us to compute these terms. We provide here, the equation satisfied by the zero order term *u*_*i*_. To this end, we replace (8) in the above equation to obtain

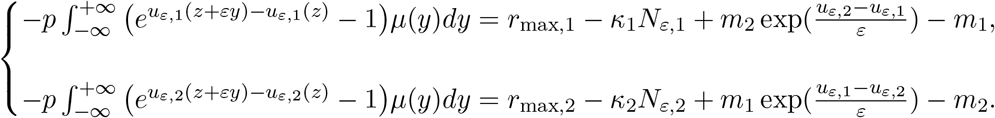

Similarly to Section A.2, the exponential terms, coming from the migration terms, suggest that when *m*_*i*_ > 0 for *i* = 1,2, as *ε* → 0, *u*_*ε*,1_ and *u*_*ε*,2_ converge to the same limit *u*. The limit u solves the following Hamilton-Jacobi equation

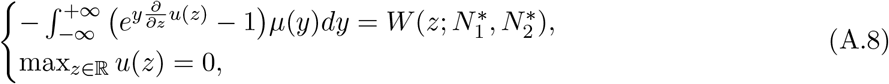

where 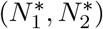 is the demographic equilibrium corresponding to the ESS and *W* is the largest eigenvalue of matrix 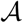 (i.e. the effective fitness introduced in Section 3.1). See [Barles et al., 2009] where the details of such computations are provided in the case of a homogeneous environment.

### B Some expressions for the case studies

#### B.1 Local densities in the dimorphic case for general and symmetric scenarios

In this section, we provide the expressions of the local densities in the dimorphic case in the adaptive dynamics framework and for the general non-symmetric scenario with *m*_*i*_ > 0. The expressions of the local densities in the symmetric case can be obtained from the same formula, using *m*_1_ = *m*_2_ =*m*, *k*_1_ = *k*_2_ = *k s*_1_ = *s*_2_ = *g r*_max,1_ = *r*_max,2_ = *r*_max_.

To this end, we first recall the values of the global densities at the ESS:

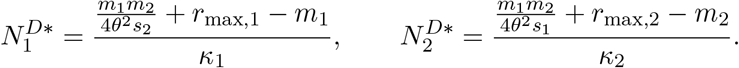

Then the local densities *v*_*k,i*_, for *k* = I, II and *i* = 1,2, are given by

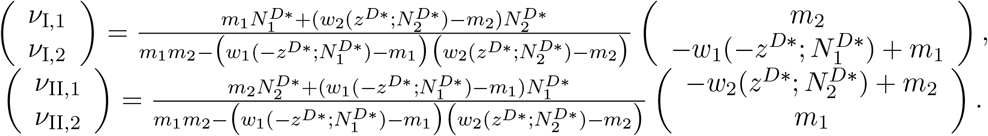

#### B.2 Condition for dimorphism in a general nonsymmetric scenario

We provide below the expressions of the constants *α*_*i*_, *β*_*i*_, and *η*_*i*_ which appear in the condition for dimorphism in Section 4.2:

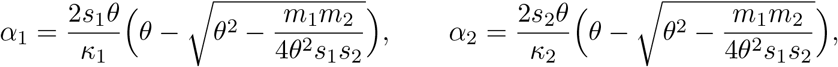

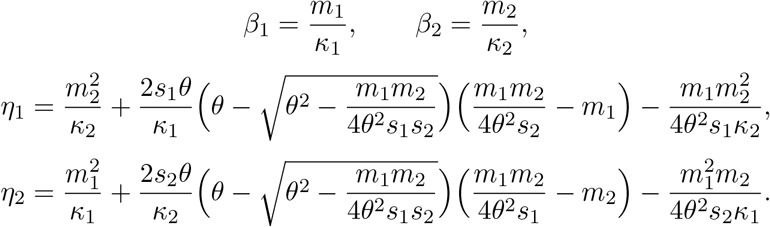

## References

Akerman, A. and Bürger, R. (2014). The consequences of gene flow for local adaptation and differentiation: a two-locus two-deme model. Journal of Mathematical Biology, 68(5):1135–1198.

Barles, G., Mirrahimi, S., and Perthame, B. (2009). Concentration in Lotka-Volterra parabolic or integral equations: a general convergence result. Methods Appl. Anal., 16(3):321–340.

Bull, J. J., Sanjuan, R., and Wilke, C. O. (2007). Theory of lethal mutagenesis for viruses. Journal of virology, 81(6):2930–2939.

Bürger, R. (2000). The Mathematical theory of selection, recombination and mutation. Wiley, New-York.

Champagnat, N., Ferrière, R., and Méléard, S. (2008). Individual-based probabilistic models of adaptive evolution and various scaling approximations, volume 59 of Progress in Probability, pages 75–114. Birkhäuser.

Christiansen, F. B. (1975). Hard and soft selection in a subdivided population. The American Naturalist, 109(965):11–16.

Day, T. (2000). Competition and the effect of spatial resource heterogeneity on evolutionary diversification. The American Naturalist, 155(6):790–803.

Débarre, F. and Gandon, S. (2010). Evolution of specialization in a spatially continuous environment. Journal of Evolutionary Biology, 23(5):1090–1099.

Débarre, F., Ronce, O., and Gandon, S. (2013). Quantifying the effects of migration and mutation on adaptation and demography in spatially heterogeneous environments. Journal of Evolutionary Biology, 26:1185–1202.

Débarre, F., Yeaman, S., and Guillaume, F. (2015). Evolution of quantitative traits under a migration-selection balance: when does skew matter? The American Naturalist, 186(37-47).

Diekmann, O., Jabin, P.-E., Mischler, S., and Perthame, B. (2005). The dynamics of adaptation: an illuminating example and a Hamilton-Jacobi approach. Th. Pop. Biol., 67(4):257–271.

Drake, J. W. and Holland, J. (1999). Mutation rates among rna viruses. Proceedings of the National Academy of Sciences of the United States of America, 96(24):13910–3.

Ducoulombier, D., Roque-Afonso, A.-M., Di Liberto, G., Penin, F., Kara, R., Richard, Y., Dussaix, E., and Féray, C. (2004). Frequent compartmentalization of hepatitis c virus variants in circulating b cells and monocytes. Hepatology, 39(3):817–825.

Fabre, C., Méléard, S., Porcher, E., Teplitsky, C., and A., R. Evolution of a structured population in a heterogeneous environment. Preprint.

García-Ramos, G. and Kirkpatrick, M. (1997). Genetic models of adaptation and gene flow in peripheral populations. Evolution, 51(1):21–28.

Gomulkiewicz, R., Holt, R. D., and Barfield, M. (1999). The effects of density dependence and immigration on local adaptation and niche evolution in a black-hole sink environment. Theoretical population biology, 55(3):283–296.

Holt, R. D. and Gaines, M. S. (1992). Analysis of adaptation in heterogeneous landscapes: implications for the evolution of fundamental niches. Evolutionary Ecology, 6(5):433–447.

Holt, R. D., Gomulkiewicz, R., and Barfield, M. (2003). The phenomenology of niche evolution via quantitative traits in a ‘black-hole’ sink. Proceedings of the Royal Society of London B: Biological Sciences, 270(1511):215–224.

Jridi, C., Martin, J.-F., Marie-Jeanne, V., Labonne, G., and Blanc, S. (2006). Distinct viral populations differentiate and evolve independently in a single perennial host plant. Journal of Virology, 80(5):2349–2357.

Kemal, K. S., Foley, B., Burger, H., Anastos, K., Minkoff, H., Kitchen, C., Philpott, S. M., Gao, W., Robison, E., Holman, S., et al. (2003). Hiv-1 in genital tract and plasma of women: compartmentalization of viral sequences, coreceptor usage, and glycosylation. Proceedings of the National Academy of Sciences, 100(22):12972–12977.

Kimura, M. (1965). A stochastic model concerning the maintenance of genetic variability in quantitative characters. Proc. Natl. Acad. Sci. USA, 54:731–736.

Kingman, J. F. C. (1978). A simple model for the balance between selection and mutation. J. Appl. Prob., 15:1–12.

Lande, R. (1975). The maintenance of genetic variability by mutation in a polygenic character with linked loci. Genetical Research, 26(3):221?235.

Lande, R. and Shannon, S. (1996). The role of genetic variation in adaptation and population persistence in a changing environment. Evolution, 50(1):434–437.

Lenormand, T. (2002). Gene flow and the limits to natural selection. Trends in ecology & evolution, 17(4):183–189.

Martin, G. and Gandon, S. (2010). Lethal mutagenesis and evolutionary epidemiology. Philosophical Transactions of the Royal Society of London B: Biological Sciences, 365(1548):1953–1963.

Meszéna, G., Czibula, I., and Geritz, S. (1997). Adaptive dynamics in a 2-patch environment: a toy model for allopatric and parapatric speciation. Journal of Biological Systems, 5(02):265–284.

Mirrahimi, S. (2011). Concentration phenomena in PDEs from biology. PhD thesis, Univeristy of Pierre et Marie Curie (Paris 6).

Mirrahimi, S. (2013). Migration and adaptation of a population between patches. Discrete and Continuous Dynamical Systems - Series B (DCDS-B), 18(3):753–768.

Mirrahimi, S. (2017). A Hamilton-Jacobi approach to characterize the evolutionary equilibria in heterogeneous environments. Math. Models Methods Appl. Sci., 27(13):2425–2460.

Mirrahimi, S., Perthame, B., and Souganidis, P. E. (2015). Time fluctuations in a population model of adaptive dynamics. Annales de l’Institut Henri Poincare (C) Analyse Non Linéaire, 32(1):41–58.

Nagylaki, T. (1978). A diffusion model for geographically structured populations. Journal of Mathematical Biology, 6(4):375–382.

Perthame, B. and Barles, G. (2008). Dirac concentrations in Lotka-Volterra parabolic PDEs. Indiana Univ. Math. J., 57(7):3275–3301.

Rice, S. H. (2004). Evolutionary theory: mathematical and conceptual foundations. Sin-auer Associates, Inc.

Ronce, O. and Kirkpatrick, M. (2001). When sources become sinks: migration meltdown in heterogeneous habitats. Evolution, 55(8):1520–1531.

Sanjuán, R., Codoñer, F. M., Moya, A., and Elena, S. F. (2004). Natural selection and the organ-specific differentiation of hiv-1 v3 hypervariable region. Evolution, 58(6):1185–1194.

Sanjuán, R., Nebot, M., Chirico, N., Mansky, L., and Belshaw, R. (2010). Viral mutation rates. Journal of Virology, 84(19):9733–9748.

Slatkin, M. (1978). Spatial patterns in the distributions of polygenic characters. Journal of Theoretical Biology, 70(2):213–228.

Szilágyi, A. and Meszéna, G. (2009). Two-patch model of spatial niche segregation. Evolutionary Ecology, 23(2):187–205.

Turelli, M. (1984). Heritable genetic variation via mutation-selection balance: Lerch’s zeta meets the abdominal bristle. Theoretical Population Biology, 25(2):138–193.

Whitlock, M. C. (2015). Modern approaches to local adaptation. The American Naturalist, 186(S1):S1–S4. PMID:26098334.

Yeaman, S. and Guillaume, F. (2009). Predicting adaptation under migration load: the role of genetic skew. Evolution, 63(11):2926–2938.

